# Ontogeny of oscillatory slow-wave and neuronal population activity in human iPSC-3D cortical circuits

**DOI:** 10.1101/2022.03.14.484311

**Authors:** Julia Izsak, Stephan Theiss, Sebastian Illes

**Affiliations:** Institute of Neuroscience and Physiology, Sahlgrenska Academy at University of Gothenburg, Sweden; Institute of Clinical Neuroscience and Medical Psychology, Medical Faculty, Heinrich Heine University, Düsseldorf, Germany; Result Medical GmbH, Düsseldorf, Germany

## Abstract

Oscillatory slow-wave activity (0.5—100 Hz) emerges during fetal human cortex development reflecting functional consequences of cellular brain ontogeny. Human induced pluripotent stem cell-derived (iPSC) neural *in vitro* models recapitulate aspects of *in vivo* cellular brain ontogeny, while neuronal mesoscale functional ontogeny is largely uncharacterized. We utilized a human iPSC-derived 3D cortical aggregate model to assess properties of emerging oscillatory slow-wave activity and its relation to synchronous neuronal population activity in cortical circuits. We reveal that oscillatory slow-wave activity (< 1 Hz), phased locked to synchronous population bursting, emerges within 14 days *in vitro* followed by consecutive stages of emerging delta (1—4 Hz), theta (4—11 Hz), beta (11—30 Hz), and gamma (30—55 Hz) oscillatory activity, accompanied by stage-specific changes in neuronal population burst pattern characteristics.

We provide a classification of neuronal mesoscale functional ontogeny stages of developing human iPSC-cortical circuits, where each stage is defined by specific oscillatory slow-wave activity and characteristic synchronous neuronal bursting patterns.

## Introduction

The mammalian brain regulates conscious and subconscious states and empowers humans and animals with sensory, motor, memory and cognitive capabilities. These brain functions fundamentally rely on electrical activity of neurons, most importantly action potentials, and adaptable synaptic transmission between neurons, mediated by smaller excitatory and inhibitory postsynaptic currents. These cellular electrical signals propagate both along axons and through neuronal and glial tissue, and can be recorded both intra- and extracellularly by electrodes. Depending on the type of electrode and its relative position to the cellular source, such electrodes can record electrical signals in the form of spikes (corresponding to action potentials) or oscillatory slow-wave activity, local field potentials (LFPs). Since Hans Berger’s first electroencephalogram (EEG) recordings in adult humans using extracranial scalp electrodes in 1929 (*1*), oscillatory slow-wave activity has been analyzed in traditional frequency bands like slow-oscillatory (< 1 Hz), delta (1—4 Hz), theta (4—11 Hz), beta (11—30 Hz), gamma (30—55 Hz) and high gamma (55—100 Hz). Sensorimotor function, speech, memory, perception, cognition and behavior—all brain information processing is accompanied by characteristic brain wave activity, reflected in the composition, power, and duration of neuronal local field potentials in these frequency bands.

Neuronal population activity and oscillatory slow-wave activity are not solely a hallmark of the adult brain, but already emerge and are detectable during fetal brain development. The first electroencephalograms of brain wave activity in full-term infants were published in 1938 by Smith et al. in a series of articles (*2–4*), and the first measurement of brain wave activity in fetal human brain was published in 1942 by Lindsey(*5*), followed by research articles describing different brain wave activity during wakefulness and sleep in pre-term infants (*6–9*). Today, properties of oscillatory activity in developing fetal human brains between gestational week 24 and birth have been well described and classified (for review see (*10, 11*)), while the early transition from immature human neurons into human neuronal circuits generating oscillatory brain activity has not yet been measured *in vivo*. Even though many different complex brain wave activity patterns emerge over time, the shift from smooth and slow brain waves with a low frequency (< 1 Hz) towards complex brain waves with higher frequency oscillatory activity (theta, beta) is a key hallmark of the functional mesoscale ontogeny of developing fetal human (for review see (*10, 11*)) and rodent postnatal neuronal circuits *in vivo* (*12, 13*).

Although rodent models have largely expanded our knowledge about how neuronal cells contribute to oscillatory slow-wave activity, these models lack human specific neural cell types (*14, 15*), limiting the translatability of results from animal models. Thus, human models are essential to better understand oscillatory activity generated from human neuronal populations. Human brain slice preparations from epileptogenic patients are suited to obtain insights into adult human neuronal circuit function (*16, 17*), but have very limited availability. Moreover, the pathological origin of such brain tissue needs to be considered when interpreting the relevance of the data for healthy human neuron and brain function.

Human pluripotent stem-cell derived neural models recapitulate the principles of cellular and morphological ontogeny of human brain development (*18*). In cortical brain organoid models, predominant delta activity has been described between 2 and 4 months *in vitro* (*19, 20*). However, these studies revealed contradictory findings regarding whether delta activity appears spontaneously (*19*), can only be elicited by epileptogenic compounds (*20*), or requires a complex neural assembloid experimental paradigm (*20*). Moreover, the timeline for the detection of delta activity in human brain organoid models is lengthy; they do not permit speedy experiments, are costly and apparently reproducibility across different research groups is challenging. Up to now, a joint systematic assessment of oscillatory slow-wave activity and neuronal population activity has not been performed in human iPSC-neural models.

Using an *in vitro* human iPSC-based 3D neural aggregate (3DNA) model on microelectrode arrays (MEAs), we provide a comprehensive description of the functional ontogeny of oscillatory slow-wave activity together with neuronal population activity patterns in developing human cortical circuits and demonstrate that late-stage human iPSC-based cortical circuits reflect characteristics of mesoscale functionality occurring in deafferented adult human cortex.

Performing weekly recordings from the same 3DNA cultures for up to three months, we reveal distinct time windows where slow-oscillatory, delta, theta, beta and gamma oscillatory activity consecutively emerged. Our data show five consecutive functional stages, where each stage has specific oscillatory slow-wave activity and characteristic synchronous neuronal bursting patterns. Our findings are repeatedly observed across different experiments and across different human iPSC-lines confirming a conserved functional ontogeny of human iPSC-derived neuronal circuits in a human 3D neural *in vitro* model resembling aspects of *in vivo* human and rodent brain development.

## Results

To assess the emergence of low frequency oscillatory activity patterns and neuronal population bursts in human cortical circuits *in vitro*, we used human iPSC derived 3D neural aggregates (3DNAs) (Fig. 1A-F). To obtain 3DNAs, we used the “dual-SMAD inhibition” protocol for neural induction of human iPSCs to obtain neural rosettes comprising neural stem cells with a dorsal telencephalic identity (fig. 1A, (*21–24*)), which were stored as cryostocks of human iPSC-cortical neural stem cells. After thawing and re-plating, neural stem cells formed neural rosettes and individual neural rosettes gave rise to 3D neural aggregates (*21–23*). 3D neural aggregates comprised cortical neurons (fig. 1C, suppl. video 1), few astrocytes (fig. 1D, suppl. video 1, (*22*)) and oligodendroglial cells (suppl. video 1, (*22, 25*)). Cortical neurons in 3D neural aggregates had mature synapses indicated by pre- and post-synaptic marker staining (fig. 1E). The cortical identity of neurons within 3DNA was confirmed by several cortical markers present in MAP-2ab^+^- and bTubIII^+^-neurons (fig. 1F). Note that cortical neurons had clear axo-dendritic orientation with MAB-2ab^+^-soma and dendrites residing within the 3DNA (fig. 1C, suppl. video 1), while bTubIII^+^-axons projected radially outside of the 3DNA (fig. 1E, F, suppl. video 1). For an extensive cellular and electrophysiological characterization of our human iPSC-3DNA experimental model see previous work (*21, 22, 25*).

**Figure 1.**
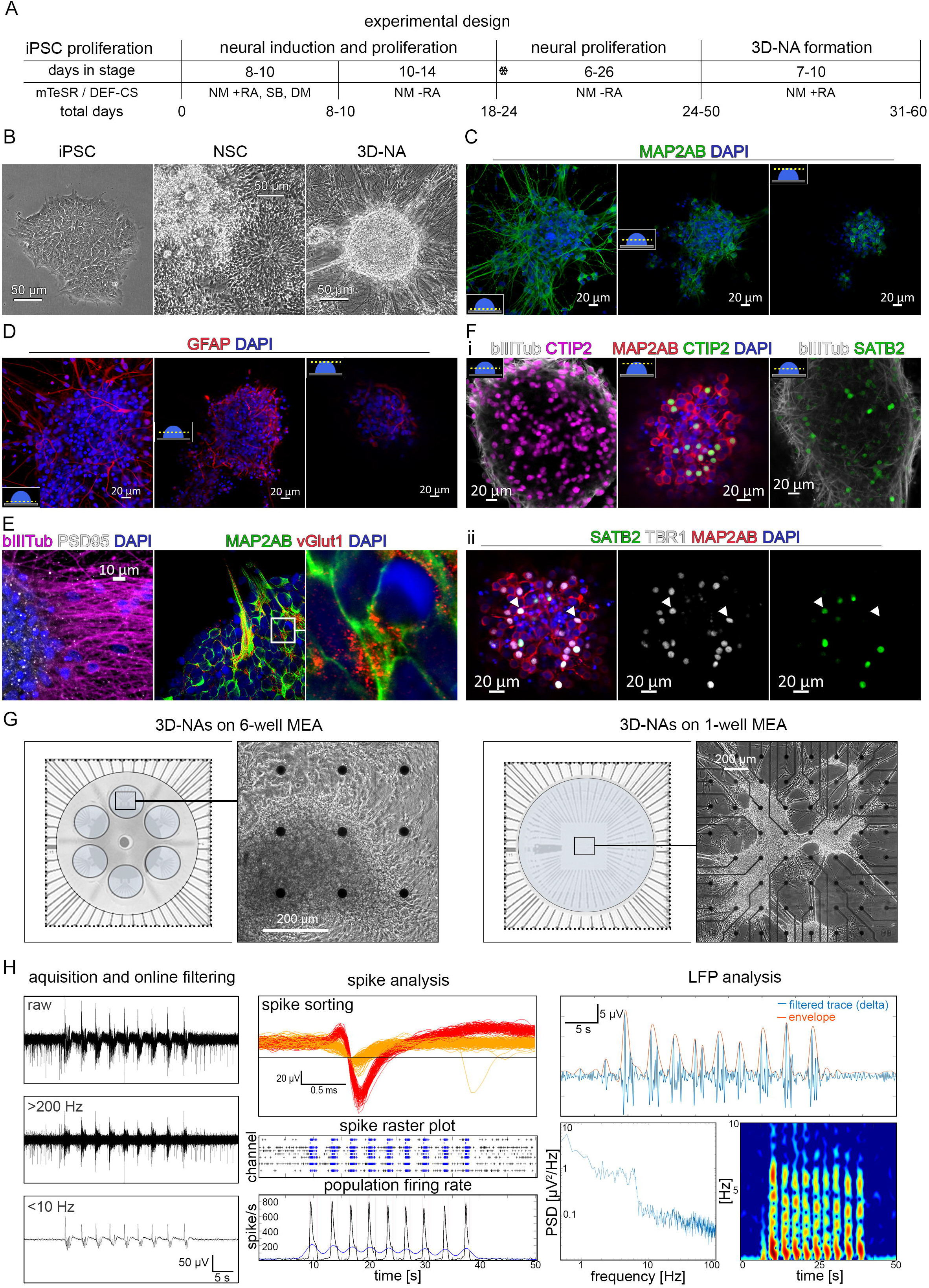
Generation and assessment of hiPSC-derived 3D-neural aggregates (3DNAs) **(A)** Schematic representation of the applied procedure, basal medium used for *in vitro* generation of hiPSC 3DNAs. **(B)** Phase-contrast images showing the morphological properties of human iPSCs, NSCs, and 3DNA. Scale bars=50 μm. **(C)** Confocal images visualizing MAP2-ab^+^-neurons at the bottom, middle and top of 3D-neural aggregate. **(D)** Confocal images visualizing GFAP^+^-astrocytes at the bottom, middle and top of 3D-neural aggregate. **(E)** Confocal images visualizing PSD95^+^-synaptic puncta and bIIItubulin^+^-neurons and vGLut1^+^-synapses in MAP2ab^+^-neurons within 3DNAs. **(F, i, ii)** Confocal images visualizing CTIP2^+^, SATB2^+^ and TBR1^+^ cortical MAP2ab^+^-neurons in 3DNAs. In **C, D, F** schematic drawings illustrate the z-level of image acquisition. **(G)** Schematic drawings, illustrating the electrode configuration in 6-well MEA and 1-well MEA, and phase contrast images showing the morphology of 3DNAs cultured on these MEAs. Scale bars=200 μm. **(H)** Image acquisition and analysis workflow from left to right: Example MEA recordings showing raw data, high-pass filtered (> 200 Hz) and low-pass filtered (< 10 Hz) data on one individual channel (2^nd^ order Butterworth filter). Exemplary spike waveform of one individual channel and exemplary spike analysis of one well of a 6-well MEA, visualizing spike raster plot and population firing diagram. Exemplary trace showing delta band filtered signal and envelope using high-order linear phase finite impulse response filter, Power Spectral Density diagram and spectrogram. Abbreviations: iPSC-induced pluripotent stem cell, NSC-neural stem cell, 3D-NA-3D-neural aggregate, NM-neural maintenance medium, DM-dorsomorphin, RA-retinoic acid, MEA-microelectrode array, LFP-local field potential, PSD-power spectral density

### Functional ontogeny of oscillatory slow-wave activity and synchronous neuronal population spike/burst activity in early human iPSC-cortical circuits

To assess whether low frequency oscillatory activity emerges and is associated with neuronal population spike/burst activity of human cortical neurons, 3D neural aggregates (50 to 60 days post-iPSC stage) were cultured either on 9-electrode 6-well MEAs or 60-electrode single-well MEAs (fig. 1G). The first set of experiments was conducted by MEA recordings every three to five days for up to 4 weeks. During data acquisition, we first assessed the emergence of neuronal spiking, single-channel-and population bursting, as well as of low frequency oscillatory activity below 10Hz by visual inspection of both high-pass (> 200 Hz) and low-pass (< 10 Hz) filtered signals (fig. 1H).

In addition, we conducted offline data processing to confirm and assess properties of low frequency oscillatory activity in both time and frequency domains. In particular, we analyzed the time domain properties shape, organization, amplitude, duration and frequency of bandpass-filtered signals in the six frequency bands slow (< 1 Hz), delta (1—4 Hz), theta (4— 11 Hz), beta (11—30 Hz), gamma (30—55 Hz) and high gamma (55—100 Hz); cf. figure 1H. While simple high- and low-pass filters (e.g. 2^nd^ order Butterworth) provided by the MEA recording software MC_Rack (Multichannel Systems (MCS)) proved sufficient for fast online visualization of both spike and slow oscillatory activity, we took care to employ high-order linear phase finite impulse response (FIR) filters for offline quantitative analysis in the separate frequency bands (cf. Methods; suppl. fig. 1). For frequency domain analyses, we calculated average power spectral densities using Welch’s method (cf. Methods) as well as power spectrograms. Eventually, we complemented our low-frequency analysis with evaluation of neuronal population bursting characteristics, including occasional spike morphology assessment (fig. 1H). In our 3DNA cultures, we regularly observed a succession of four different activity states, defined by both specific slow-wave activity and concomitant spike/burst patterns that we categorized as “type A” to “type C”.

As described previously, human cortical neurons within the human 3DNA cultures establish synchronously spiking neuronal networks within 30 days after plating on MEAs, where the first spontaneous synchronous neuronal population bursts appear within 10—14 days *in vitro*(*23*). Here, with the appearance of the first population bursts (10—14 div), we observed low frequency oscillatory activity (0.5—1 Hz) that was barely visually detectable during online assessment (fig. 2A, column “Type A”). Spectrograms confirmed the presence of spontaneous low frequency oscillatory activity, which occurred during spontaneous synchronous population bursts (fig. 2A, type A). This indicates that low frequency oscillatory activity (< 1 Hz) is already associated with the early emergence of population bursts. We termed this early spontaneous neuronal network activity “Type A”, characterized by slow-wave oscillatory activity with duration less than 1 second and low amplitude, where population bursts are phase-locked to the rising flank of the slow-wave.

**Figure 2.**
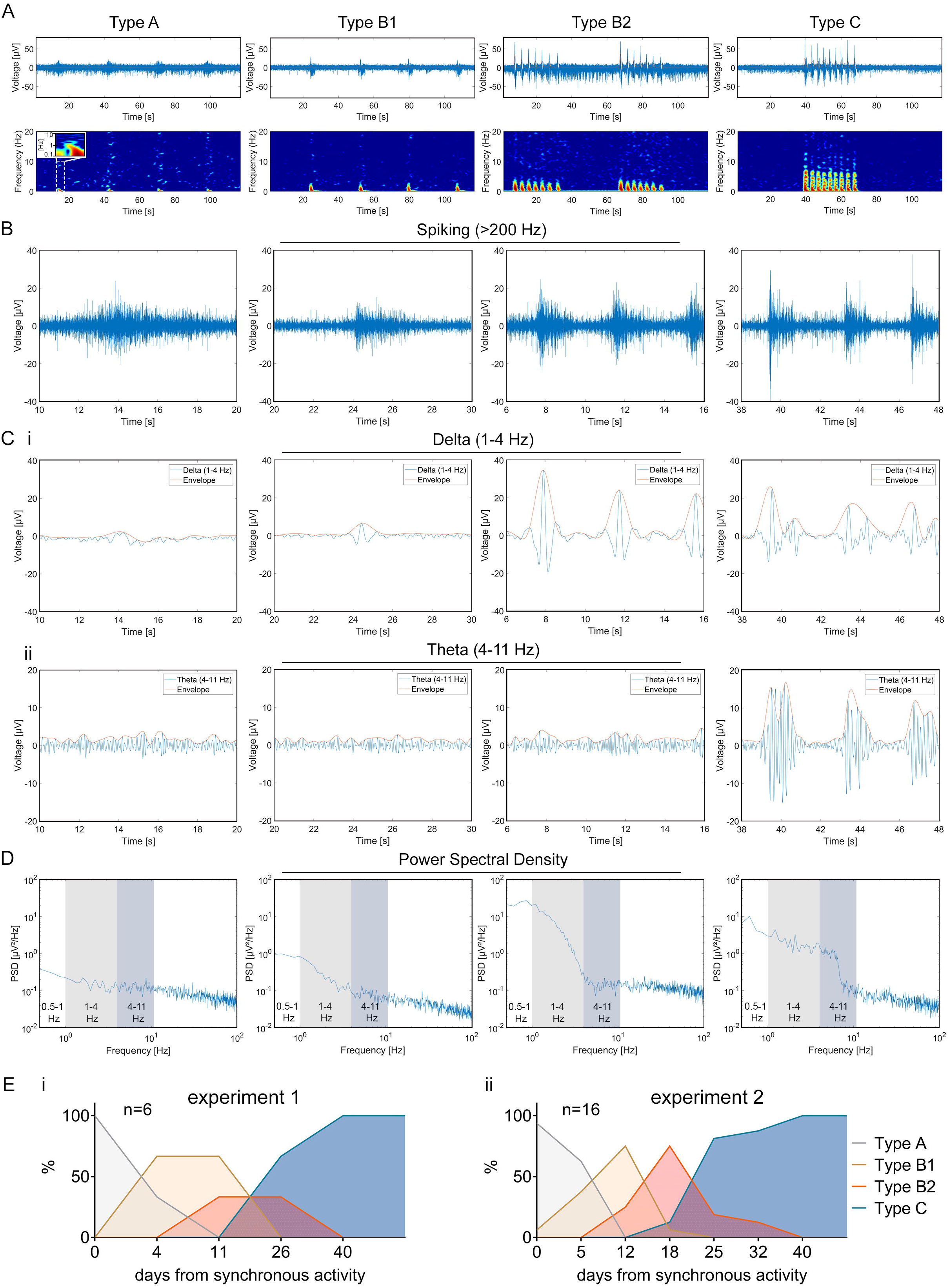
Early ontogeny of synchronous bursting and oscillatory slow-wave activity. **(A)** Representative examples of 2-minute raw activity traces (blue) and superimposed delta envelope (orange) of different neuronal activity states (from type A to C, left to right) and their corresponding spectrograms. Zoomed-in 10 s window traces showing **(B)** the filtered spiking signal, **(C, i)** the filtered delta signal and envelope, and **(C, ii)** the filtered theta signal and envelope. **(D)** Power Spectral Density graphs corresponding to the different neuronal activity states, where the delta (grey) and theta (blue) frequency bands are highlighted. **(E)** Diagrams show the percentage of cultures exhibiting type A, B1, B2, or type C network activity characteristics over time after the first detection of synchronous spiking activity (day 0) in experiment 1 **(i)** and 2 **(ii)**.

After the appearance of type A network activity, we observed three other network activity types emerging within the next two weeks (14—30 div). In the following, we introduce human network activity types B1, B2 and C, and their distinct characteristics (fig. 2). Type B1 network activity is defined by an increased number of population bursts (PBs), higher degree of neuronal network synchrony, slow oscillatory activity with increased amplitude and the emergence of delta band activity (1—4 Hz) (fig. 2, type B1). Type B2 network activity is characterized by the appearance of population super bursts (PSBs) composed of several consecutive PBs nested on slow-wave oscillatory delta band activity (fig 2, type B2). Type C network activity is characterized by longer PSBs composed of more consecutive PBs nested on slow-wave oscillatory delta band activity and prominent theta band (4—11 Hz) oscillatory activity (fig 2, type C).

Next, we asked if the different network activity types systematically appeared in a specific order during the early development of human cortical *in vitro* circuits. For this purpose, we assessed six individual 3DNA cultures from one and 16 individual 3DNA cultures from another experiment. We plotted the percentage of cultures showing type A, B1, B2, or type C network activity characteristics over time after the first detection of low-power slow oscillatory activity and associated population bursting (type A network activity). As shown in figure 2E, all 22 individual human neuronal circuits from two independent experiments developed from type A to B1, from type B1 to B2 and subsequently from type B2 to C.

These data provide evidence that in early synchronously spiking human iPSC derived cortical circuits an ontogeny of distinct early functional stages occurs and is characterized by a time-dependent shift in slow oscillatory activity from low (< 1 Hz) towards higher (delta, theta) frequencies. Network activity types A, B1 and B2 occur transiently, before a temporal plateau network stage (type C) develops within 40 days after the first onset of synchronous activity.

We repeated these experiments with 3DNA cultures obtained from three other human iPSC lines in 17 independent experiments, where each experiment comprised several individual 3DNA cultures (suppl. figure 2). By using the data from 124 individual 3DNA cultures, we confirmed that the functional development trajectory given by transitions from type A → B1 → B2 → C occurred within 25—40 days after the first onset of synchronous spiking (suppl. fig 2B).

While the transition time points from one oscillatory activity type to the next showed temporal variability between individual 3DNA cultures and experiments, we reproducibly detected a clear sequential development trajectory A → B1 → B2 → C: emergence of distinct low frequency oscillatory activity types accompanied by distinct neuronal population burst pattern characteristics. Our data demonstrate that the formation of synchronous neuronal networks is not the endpoint of network maturation, which indicates that beyond functional network formation, additional *in vitro* conserved sequential cellular and synaptic neurodevelopmental processes likely occur that are reflected in the consecutive emergence of slow oscillatory activity types and population burst patterns in early developing human cortical circuits.

### Origin and properties of oscillatory slow-wave activity in human iPSC-derived 3DNA cultures

Next, we showed that suppression of fast glutamatergic transmission by chemical inhibition of NMDA- and AMPA-receptors resulted in the elimination of synchronous neuronal population bursting and slow oscillatory activity, while uncorrelated neuronal spiking detected by several electrodes was preserved (suppl. video 2). Elimination of synchronous neuronal population bursts and oscillatory slow-wave activity could also be achieved by blocking voltage-gated sodium channels after tetrodotoxin (TTX) application (suppl. video 2). Under TTX conditions, no uncorrelated neuronal spiking could be detected (suppl. video 2). In addition, no slow oscillatory activity could be detected in early 3DNA cultures (within 7 div), which only exhibited uncorrelated neuronal spiking activity (suppl. fig. 3). These experiments emphasize that oscillatory slow-wave activity recorded in early developing human *in vitro* cortical circuits depends on neuronal excitability and excitatory synaptic communication— both also prerequisites of synchronous neuronal population activity.

Local field potentials recorded by electrodes *in vitro* or *in vivo* are considered to reflect voltage fluctuations, summed up over a ball of ~ 200 μm diameter, that originate from dipole currents during synchronous activity of oriented neurons and then propagate through neuronal tissue (*26*). Since cortical neurons in our 3DNA showed a clear axo-dendritic orientation (fig. 1, fig 3B and suppl. video1), we assessed if the detection of slow-wave oscillatory activity required that electrodes were covered by clusters of neurons within 3DNAs (fig. 3A, suppl. video 1). To this end, we examined 291 electrodes detecting oscillatory slow-wave activity for simultaneous 3DNA coverage as a possible prerequisite.

**Figure 3.**
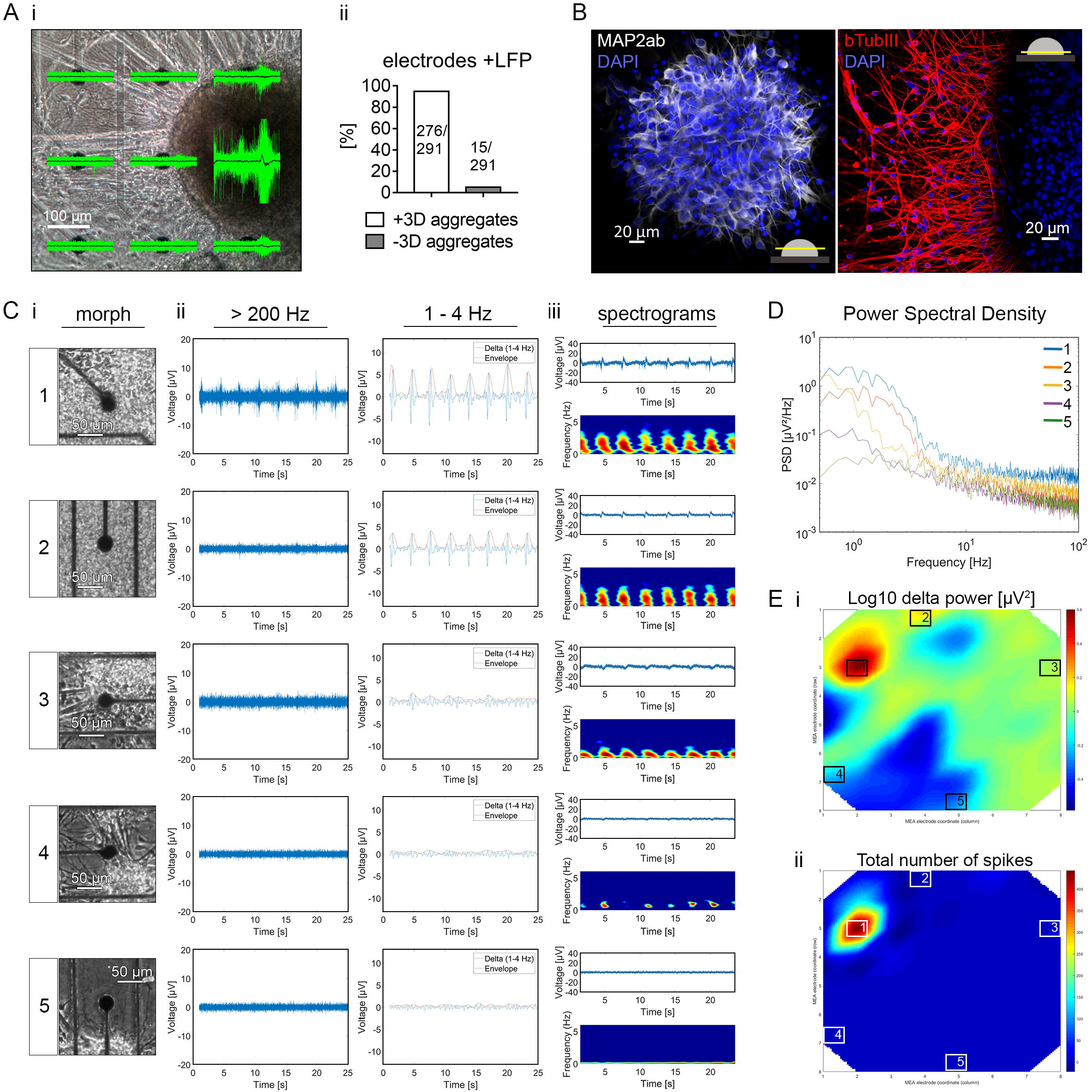
Oscillatory slow-wave activity originates from neurons with a defined axo-dendritic orientation localized within 3DNAs. **(A, i)** Phase-contrast image, showing the morphology of a 3D-neural aggregate 14 days after neural aggregate isolation and cultivation on the 9 electrodes of a 6-well MEA. The superimposed green traces visualize spiking and black traces visualize the delta signal recorded from the culture. **(A, ii)** Diagram shows the percentage of electrodes recording local field potentials when covered (white) or not covered (grey) by 3DNAs. The number of analyzed electrodes is given. **(B)** Confocal images visualizing MAP2ab^+^-neurons in the middle, and bIIItubulin^+^-neurons at the bottom of 3DNAs, stained 19 days after seeding 3DNAs on coverslips. **(C, i)** Phase-contrast images, showing the morphology of 3D-neural aggregate cultures on 5 highlighted electrodes of a 60-electrode MEA, 21 days after seeding. Numbered boxes in **(E)** mark the highlighted electrodes, and corresponding recorded activity is shown in **(C, ii, iii, D, E)**. **(C, ii)** Filtered spiking signal, delta band filtered signal and envelope, **(C, iii)** raw activity traces (blue) and superimposed delta envelope (orange) and their corresponding spectrograms. **(D)** Power Spectral Density graphs corresponding to the highlighted electrodes, **(E)** Spatial heat map showing detected **(i)** delta power and **(ii)** spiking activity on the 8 × 8 microelectrode array.

Indeed, roughly 95% of all electrodes detecting slow-wave oscillatory activity were covered by 3DNAs (276 out of 291 electrodes, 94,8%, fig. 3A), consistent with the concept that 3-dimensional neuronal clusters with axo-dendritic orientation strongly support slow-wave oscillatory activity that most likely cannot be detected from individual neurons or axonal processes.

Interestingly however, roughly 5% of electrodes detecting low frequency oscillatory activity were not directly covered but rather in close vicinity to 3DNAs (15 out of 291 electrodes, 5.15%, fig. 3A, ii), indicating that electrodes can detect oscillatory slow-wave activity even at some lateral distance from the neuronal signal source. While axonal propagation of action potentials (APs) preserves AP waveforms over long distances, electrical signals propagating through a liquid or solid medium are attenuated in a frequency dependent manner: for low-frequency electrical signals (e.g. delta waves, 1—4 Hz), attenuation is much lower than for high frequencies as e.g. in spikes (> 1000 Hz), which is consistent with our observation of delta band oscillatory activity on electrodes distant from 3DNAs.

We therefore surmise that low-frequency oscillatory activity may more sensitively detect the presence of neuronal activity than neuronal spiking. To test this hypothesis, we conducted an experiment where twenty to thirty 3DNAs were distributed and cultured on an 8 × 8 microelectrode array. On these MEAs with a 1.84 × 1.84 mm^2^ recording area, each 30 μm diameter electrode has a center-to-center distance of 200 μm to its neighboring electrodes. In recordings of spatially distributed 21 div human iPSC-derived 3DNA cultures, we observed that only few electrodes detected spiking activity—in the exemplary recording shown in figure 3C in the form of population super bursting—, while considerably more electrodes detected low frequency oscillatory delta band activity (1—4 Hz) (fig. 3C, ii). The sole assessment of spike activity by e.g. spatial heat maps provided the impression that only a minor fraction of cultured 3DNAs contained electrophysiologically active neurons and that the spatially distributed individual 3DNAs were not functionally interconnected (fig. 3E ii). In contrast, spatial heat maps of slow-wave oscillatory activity revealed that many individual 3DNAs contained electrophysiologically active neurons, and that neurons within individual 3DNAs were functionally interconnected as evidenced by spontaneous synchronous slow oscillatory activity detected in spatially distributed individual 3DNAs (fig. 3E i). Thus, measurements of slow-wave oscillatory activity provide complementary information about neuronal activity and interconnectivity of spatially distributed human 3D neural aggregates.

### Functional ontogeny of oscillatory slow-wave activity and synchronous neuronal population spike/burst activity in late-stage human iPSC-cortical circuits

In a second set of experiments, we assessed if the so far observed slow-wave oscillatory activity and population burst patterns persisted in long-term cultures of 3DNAs, and if additional low frequency oscillatory activity and population burst patterns emerged over time. For this purpose, we cultured 3DNAs on 9-electrode arrays (6-well MEAs) for up to three months and applied the same assessment and analysis paradigms as used in the first set of experiments.

3DNA cultures consecutively developed into the types A → B1 → B2 →C of synchronously active neuronal networks (fig. 4, first column) confirming once more the results from our first set of experiments.

**Figure 4.**
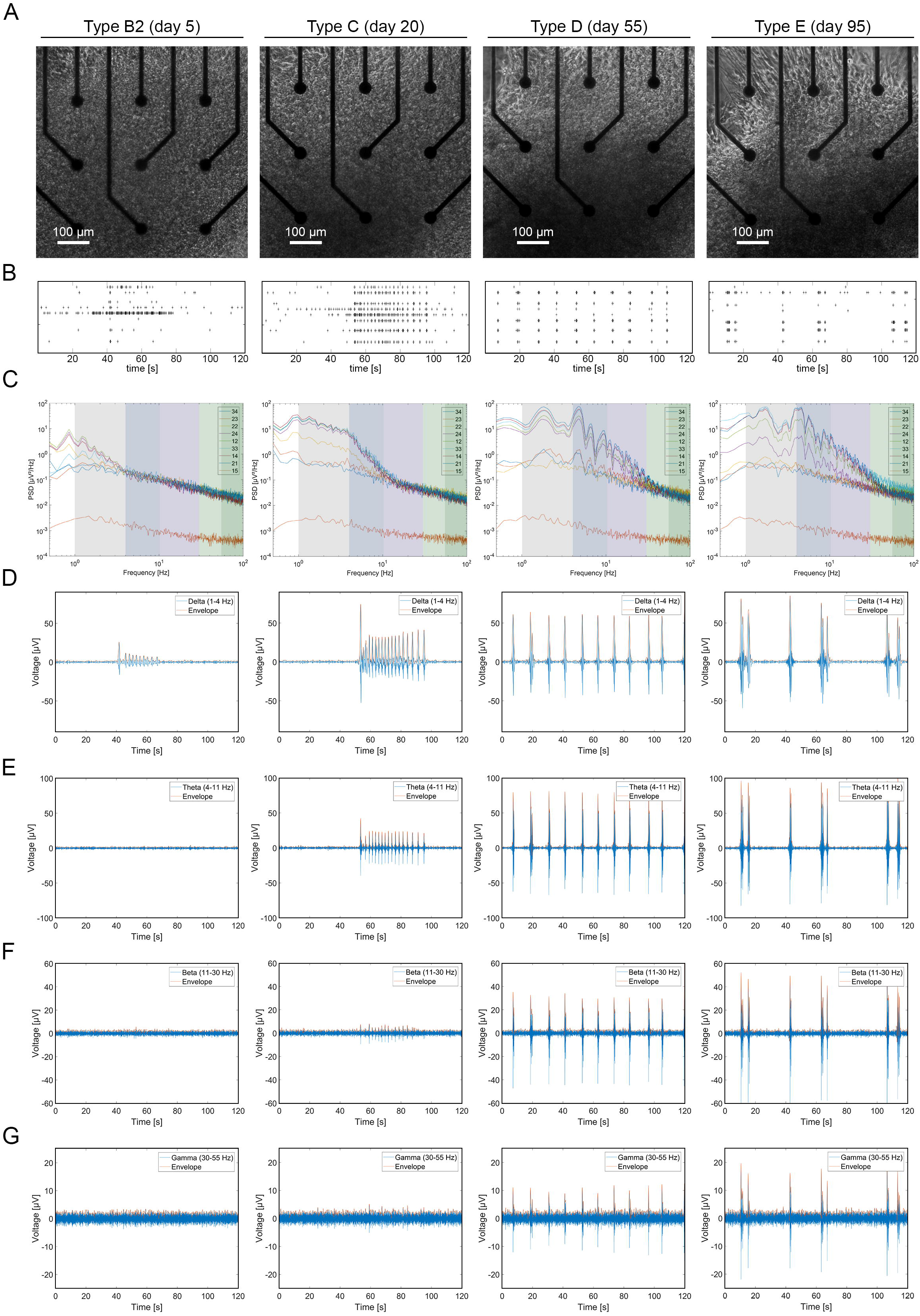
Late ontogeny of synchronous bursting and oscillatory slow-wave activity. **(A)** Phase-contrast images, showing the morphology of a 3D-neural aggregate culture on the 9 electrodes of a 6-well MEA over the time course of 120 days. Indicated days represent the days after the first onset of synchronous neuronal activity corresponding to the different activity stages from Type B2 to E. Corresponding **(B)** spike raster plots and **(C)** Power Spectral Density graphs, where the different frequency ranges are highlighted by coloured boxes. Representative one channel filtered traces showing **(D)** delta trace and envelope, **(E)** theta trace and envelope, **(F)** beta trace and envelope and **(G)** gamma trace and envelope representative for the different activity states.

At later times, however, we observed another neuronal network activity pattern (type D) where prominent beta band oscillatory activity (11—30 Hz) emerged and PSB was absent and PBs appear as single events (starting around 40 to 50 days after the onset of synchronous activity (fig 4, third column).

Even longer cultivation times revealed the emergence of yet another type (type E), where PBs either occurred as groups of three to four PB or as single events. A key hallmark of type E neuronal network activity was the emergence of gamma band (30—55 Hz) oscillatory activity (fig 4, fourth column). In all experiments and all cultures, we observed that the peak amplitudes of all oscillatory activity envelopes showed a progressive increase over the time course of three months (fig. 4D-G).

We also noticed that the structure of individual population bursts and the wave form of slow (< 1) and delta band (1—4 Hz) oscillatory activity changed over time (fig. 5). In detail, PBs during early neuronal circuits (type A—C) is a single PB nested on a single smooth wave with a frequency around 1 Hz. In late neuronal circuits (type D and E), the duration of an individual PB was longer (1.5 seconds), and the PBs comprised 6 to 8 sub-PBs (fig. 5). Interestingly, these PB characteristics remained stable for an additional month of cultivation. Furthermore, we could not observe other types of neuronal network activity patterns in long term cultures performed systematically over a time of three months, and occasionally up to four months. We repeated the experiment with another human iPSC cell line and observed the emergence of identical developmental stages within comparable timelines (suppl. fig.4). Thus, we conclude that type E network activity represents a final functional stage of *in vitro* human cortical circuit activity.

**Figure 5.**
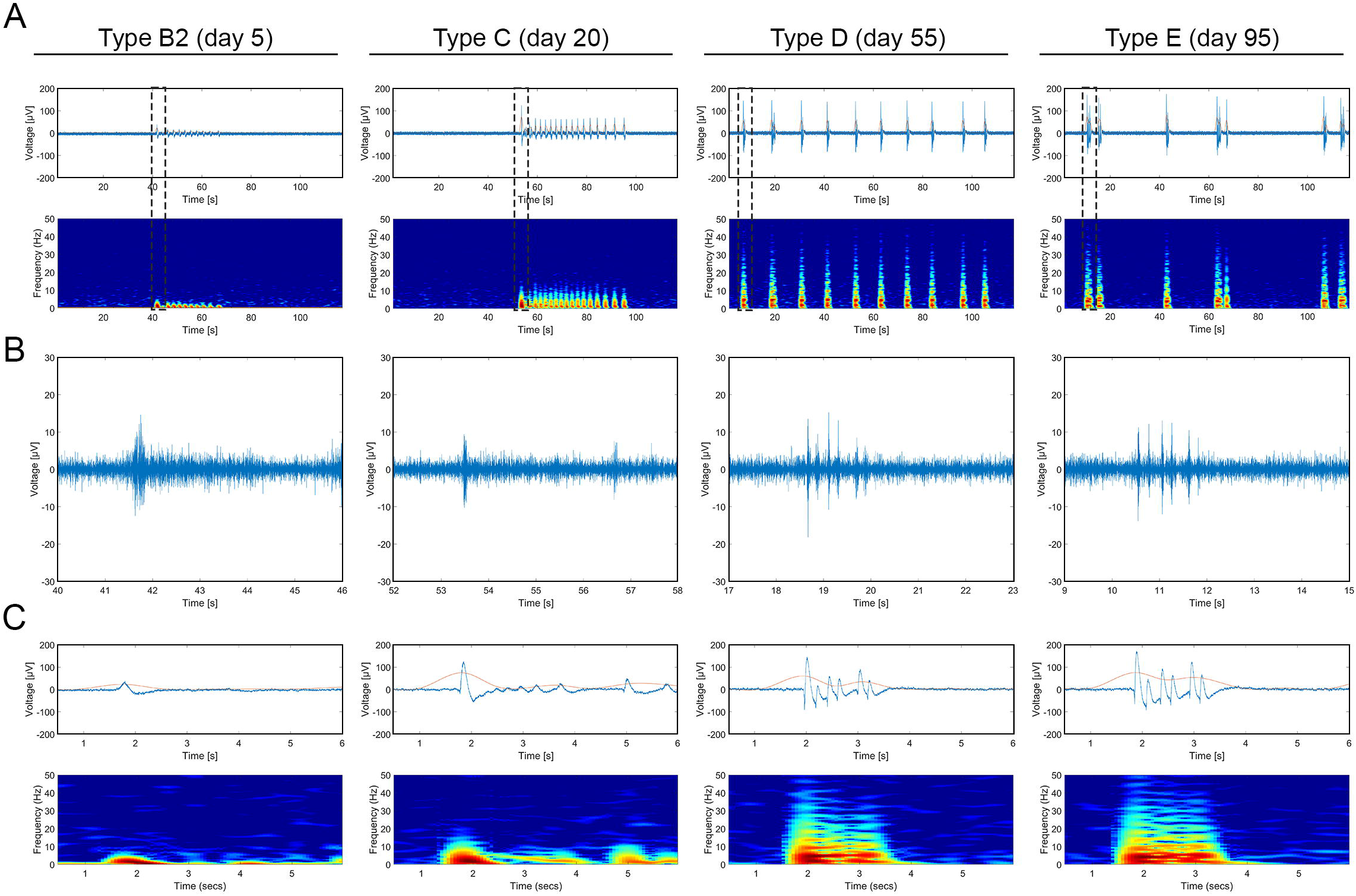
Transition of smooth oscillatory slow-wave activity to nested oscillatory activity during human neuronal circuit development in 3DNAs. **(A)** Representative examples of 2-minute raw activity traces (blue) and superimposed delta envelopes (orange) for different neuronal activity states (from type B2 to E, left to right) and their corresponding spectrograms. The marked rectangles are shown below as zoomed-in 6 s window traces showing **(B)** the filtered spiking signal, **(C)** raw activity traces (blue) and superimposed delta envelopes (orange) and their corresponding spectrograms of different neuronal activity states (from type B2 to E, left to right).

To summarize, we demonstrate a functional ontogeny in human 3D aggregates of brain cells which consists of five consecutive functional developmental stages, where each stage has specific oscillatory slow-wave activity and characteristic synchronous neuronal bursting patterns (fig.6).

**Figure 6.**
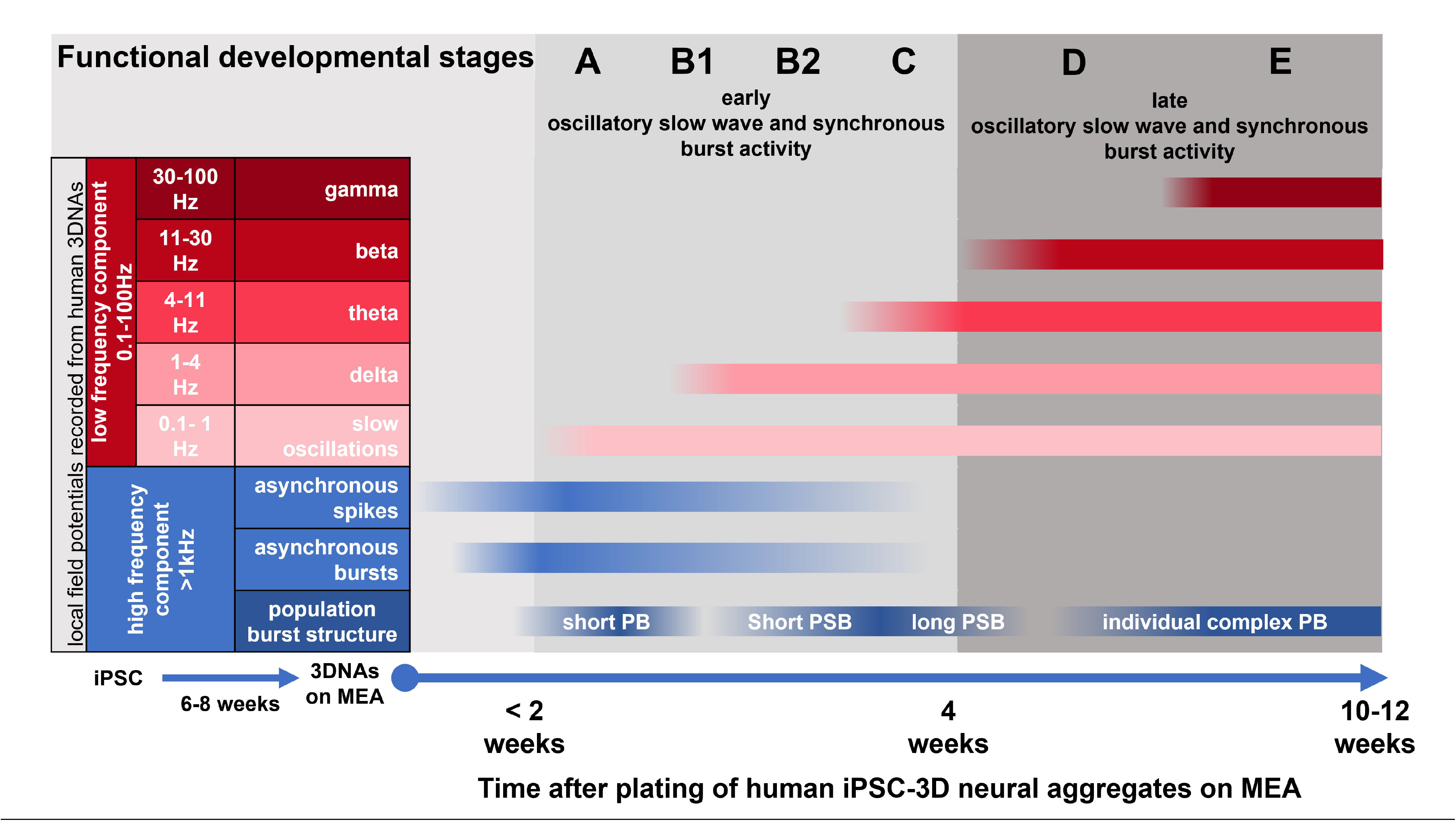
Schematic overview of functional development stages. Human iPSC-derived 3D neural aggregates (3DNAs) generate local field potentials composed of a high-frequency component (>1kHz, neuronal spiking) and a low frequency component (oscillatory slow-wave activity). Schematic illustration summarize the functional ontogeny in human 3D aggregates of brain cells which consists of five consecutive functional developmental stages, where each stage has specific oscillatory slow-wave activity and characteristic synchronous neuronal bursting patterns

## Discussion

Progress in understanding physiological and pathological oscillatory human brain activity has been hampered by limited access to primary human tissue and limited translatability of animal models. Mechanisms involved in the emergence of oscillatory activity in fetal human neuronal networks cannot be studied during very early human brain development stages *in vivo*, and scalp electrodes applied on pre-term babies do not have sufficient spatial resolution to capture neuronal spiking or bursting activity. Moreover, animal models lack human specific neural cells (*14, 15*). Thus, an accessible human neuronal *in vitro* model, in which the emergence and properties of neuronal population bursting and oscillatory slow-wave activity can be simultaneously studied, represents an opportunity to overcome current limitations. The accessibility of human iPSC-derived neural *in vitro* models will uniquely enable intervention studies and provide novel insights about human neuronal circuit function. In this light, we believe assessing the emergence and properties of oscillatory activity recorded from hiPSC-neuronal circuits *in vitro* will help to better understand human brain function.

Our experimental paradigm of adherently growing 3D neural aggregates has been utilized to assess the development and properties of synchronous mouse (*27*) and human pluripotent stem cell-derived neuronal population bursting (*23*), to evaluate effects of clinically relevant compounds on human neuronal network function (*28*), and to provide insights into the cellular neural ontogeny (*29*) and corticogenesis (*30*). For instance, Edri *et al.* presented the cellular ontogeny of neuroepithelial cells and radial glia cells within neural rosettes (*29*). We and others (*29*), have provided a detailed description about how individual neural rosettes give rise to adherently growing 3D neural aggregates (*23, 29*). Thus, our experimental paradigm of 3D neural aggregates represents a fast, robust and cost-efficient 3D human neural *in vitro* model to assess specific aspects of cellular and functional human brain ontogeny *in vitro*.

While brain organoid models have recently gained popularity in science, this experimental paradigm has several drawbacks, among which there are lengthy and costly experiments, high experimental variability, insufficient and unphysiologically long neuronal maturation and high degree of cell stress (for a recent review see (*31*)). In this work, we aimed to evaluate if human iPSC-derived 3D neural aggregates provide insights into the emergence of oscillatory slow-wave activity in developing human iPSC-cortical circuits.

The use of hiPSC-derived neural *in vitro* models to assess oscillatory activity of human neurons *in vitro* is still in its infancy. As of today, there is only one systematic published study (*19*) about the emergence and properties of delta band oscillations and changes of population burst characteristics in developing human neurons, viz. brain organoids. In the following sections, we will compare and discuss currently available data about the emergence of oscillatory slow-wave activity in brain organoids with the data obtained from 3DNAs presented in this work.

### Developmental and morphological properties of 3DNA and brain organoids

Similar to human brain organoids, 3D neural aggregates comprise neurons (fig. 1C, E, F, suppl. video 1), few astrocytes (fig. 1D, suppl. video 1) and oligodendroglial cells (suppl. video 1, REF), as well as more immature neural progenitor and neural stem cells (*22*). While human brain organoids are cultured free-floating, and often consist of several neural rosettes, 3D neural aggregates are cultured adherently. Each 3DNA derived from an individual neural rosette that exhibits apical-basal orientation, shows interkinetic nuclear migration and neuronal differentiation at the apical side (*23*) similar to neural rosettes within free-floating brain organoids (*19*). Interestingly, neither neurons within brain organoids, nor neurons within 3D neural aggregates are organized in layers. In detail, MAP-2ab^+^-neurons do express cortical-layer specific transcription factors (layer I to layer VI), however in the absence of cytoarchitecture. Even though brain organoids have been cultured for months up to years, only neural stem/progenitor cells within the proliferation zone show a cytoarchitecture resembling the layer specific transcription factor expression profile seen *in vivo*. Nevertheless, complex neuronal population bursting and oscillatory slow-wave activity emerges in both human 3D neural *in vitro* models (*19, 32*), which demonstrates that a layered organization of neurons is not a prerequisite for complex neuronal population functionality. In addition, primary cultures of mouse neurons, where a layered organization of neurons is also absent, show robust oscillatory activity within the delta, theta, beta and gamma bands (*33, 34*). When combining the data presented here with the data published by others, it becomes evident that layered organization of neurons is not crucial for their ability to generate slow-wave oscillatory activity, and may not even be crucial for brain function, as discussed by others (*35, 36*). As our data demonstrate, oscillatory activity is primarily detectable in 3DNAs where neurons have axo-dendritic orientation, which seems to be a more important morphological prerequisite than layered organization.

### Mesoscale functional ontogeny of neurons in 3DNA and brain organoids

In this section, we aim to compare and discuss the timelines for the emergence and properties of population burst characteristics and oscillatory activity reported for neurons in brain organoids and for neurons in our 3DNA experimental paradigm presented here. A particular challenge for this comparison is that the protocols for brain organoid and 3DNA formation are different with respect to timelines for neural induction, NSC expansion, as well as organoid or 3DNA formation. Since we are interested to compare the timelines and properties of mesoscale neuronal functionality in both 3D human neural *in vitro* models, we believe it is reasonable to start this comparison based on time points when brain organoids and 3DNA, respectively, were placed on the MEA. Even though there is currently only one study by Trujillo et al. (*19*) systematically describing the developmental emergence of oscillatory activity in brain organoids, combining this data set with our data presented here allows the discovery of remarkable similarities and differences.

The timelines from iPSC to brain organoid/3DNA formation, respectively, and placement on MEAs are 7—8 weeks (3DNA) and 10 weeks (brain organoids). It is important to note that brain organoids have been pre-matured for 6 weeks before placed on MEAs.

Within the first week after plating, neurons within 3DNA cultures show asynchronous activity, and they develop into synchronous neuronal networks within additional one to two weeks, as presented here and reported elsewhere (*21–23, 28*). As described by Trujillo et al. (*19*), neurons within adherently growing brain organoids show asynchronous activity one day after plating of 6-week old brain organoids on MEAs. Within additional two weeks, also neurons in brain organoids exhibit synchronous activity(*19*). Thus, neurons in human 3DNA and brain organoids exhibit nearly identical development timelines from electrophysiologically functional neurons with uncorrelated activity to the formation of functionally interconnected and synchronously active neuronal populations.

The formation of synchronously active neuronal networks has often been considered as a mature developmental endpoint (*23, 37–39*). However, as presented here and also described by Trujillo et al., this only reflects the first stage of neuronal mesoscale functionality, which is followed by additional changes in neuronal population burst and slow-wave oscillatory activity characteristics. In detail, 3DNA cultures show consecutive emergence of oscillatory slow-wave (onset within 2 weeks), delta (onset within 2—3 weeks), theta (onset within 3—4 weeks), beta (onset within 4—6 weeks) and gamma/high gamma (onset within 6—8 weeks) activity, which are only detectable during synchronous neuronal activity. Trujillo et al. only characterized the emergence of delta activity and presented data about higher frequency bands (200—400 Hz), which are classified in the field rather as ripple activity (*40*) than gamma activity (*40*). As for the 3DNA model, delta activity (1—4 Hz) in brain organoids occurs exclusively during synchronous population bursts.

Another similarity between neurons in brain organoids and 3DNAs lies in characteristic changes of population bursts over time. In both 3D *in vitro* model systems, early PBs are < 1 sec long, appear as single synchronous events that progressively change over time into complex 1.5 to 2 seconds long PBs comprised of several sub-PBs. Since PBs are phase-locked to delta oscillations, changes in PB characteristics are also reflected in delta band wave forms, i.e. over time a transition of one delta wave with a wavelength < 1 second into 4 to 7 waves within a 2-second time window occurs. However, those characteristic changes in PB and delta band activity occur on different time scales: in 3DNAs within 4—6 weeks, and in brain organoids within 6 months (*19*). Along this line, delta power recorded from brain organoids and 3DNA cultures increases over time, presumably due to increased numbers of neurons participating in generating this oscillatory activity.

In addition, we demonstrated that population superbursting (PSB) is a neuronal population activity pattern emerging in a distinct stage during the development of early 3DNA cultures. Trujillo *et al.* (*19*) presented a spike raster plot showing clear presence of PSB in 10 months old brain organoids (suppl. figure 10); it is, however, so far unknown, if this is a transient activity pattern in brain organoids.

To summarize, the mesoscale functional ontogeny of neurons within brain organoids and 3DNAs share comparable aspects of developmental trajectories regarding (i) neuronal circuit formation, (ii) emergence and (iii) changes of population bursts and delta band activity, which indicates that conserved cellular developmental mechanisms may occur in both human 3D *in vitro* models. In addition, human neurons in 3DNAs consecutively generate spontaneous theta, beta and gamma band oscillations during distinct developmental stages, which has not been reported for neurons in brain organoid models yet. Interestingly, the timelines for the transition from asynchronous firing to synchronously active neurons in brain organoids and 3DNA are identical, while the timelines for later mesoscale functional ontogeny of neuronal populations are substantially shorter for 3DNAs.

### Slow-wave oscillatory activity of human neurons in vitro and in vivo

An important aspect to consider when comparing slow-wave oscillatory activity of human neurons *in vitro* and *in vivo* is that human brain organoids and 3DNAs are input-deprived. In principle, *in vitro* oscillatory activity could reflect activity properties observed in (i) isolated human brain slice preparations, (ii) deep NREM sleep EEG in adults, when cortical activity is largely deprived of and not modulated by sensory input from other brain regions(*41*), (iii) pre-term baby EEG at 24 to ~ 28 gestational weeks (gw), where sensory input is not fully established (*10*), or (iv) pre-term baby EEG at 30-31 gw until birth during quiescent stages (*10*) equivalent to non-REM sleep in adults.

Interestingly, recordings from pre-term infants show that within a time-period of 7 weeks (between gestational weeks 24 and 31) the transition of very slow oscillatory activity (< 1 Hz) to alpha-beta band oscillations (8—30 Hz) nested to delta band activity (1—2 Hz) is a general feature of human brain development (*10, 11*). Interestingly, we also observed a transition from low-to-high frequency oscillatory activity in our human 3D brain cell cultures occurring within weeks, and not months(*19*).

To summarize, it is tempting to discuss similarities and differences of emergence and properties of oscillatory slow-wave activity of human iPSC derived neurons *in vitro* compared to those recorded from human neurons in isolated brain slice preparations or recorded from human fetal and adult brain, but further studies are needed to reach a conclusion.

### Future outlook

We demonstrate that human 3D neural aggregate cultures represent a fast and robust human *in vitro* model for assessing the functional ontogeny of low frequency oscillatory activity and neuronal population activity during human neuronal circuit formation.

Functional ontogeny of oscillatory slow-wave activity and synchronous neuronal bursting patterns represents a novel concept with a high potential to reveal abnormal human neuronal circuit development and function when applied on human iPSC models for human brain diseases and disorders. The described functional ontogeny in brain organoids and human 3D neural aggregate cultures presented here pave the way to identify prospective abnormal development and function of oscillatory slow-wave activity in human iPSC-based CNS disease models.

## Methods

### Generation of human iPSC-derived 3D neural aggregates (3DNAs)

The method is extensively described elsewhere (*23*). Shortly, hiPSC lines (C1, C2, C3) and the commercial human iPSC-line Chipsc4 (Takara, Sweden) were cultured and differentiated into cortical NSCs as described elsewhere (*23*). Within 10–14 days, hiPSC NSCs formed 3DNAs(*23*) and 3DNAs were manually transferred on MEAs and cells were kept in BrainPhys medium with supplements.

### Immunocytochemistry and confocal imaging

For the immunocytochemical characterization of human 3DNAs, human 3DNAs were seeded and cultured on PLO/Laminin-coated 96-well plates (Greiner) and cultured in BrainPhys media with supplements (*23*). The cultures were then washed in phosphate-buffered saline (PBS), pH 7.2, and fixed for 20◻min in 4% paraformaldehyde at room temperature. After fixation, the cells were incubated with 1% bovine serum albumin for 30◻min. Primary antibodies binding to neuronal (bIIITub, MAP2ab, TBR1, CTIP2, SATB2, and Parvalbumin-PV), astrocytic (GFAP), and synaptic (PSD95, VGlut1) structures were diluted in blocking solution with 0.025% Triton-X and were applied at 4◻°C overnight. After washing in PBS, appropriate secondary antibodies together with DAPI nuclear counterstaining were applied for 2◻h at room temperature. Images were collected with a confocal-laser scanning microscope (LSM 700 Zeiss).

### MEA recordings and analysis

3DNAs were plated on either 6-well (3 × 3 electrodes) or single-well (8 × 8 electrodes) MEAs coated with PDL and laminin as previously described(*23*). Before start of recording, 3DNA cultures were left to equilibrate for ten minutes. For each culture condition, we performed five consecutive 2-minute long recordings for each 3DNA culture in culture medium.

Raw signals were recorded with a MEA2100 system (Multichannel Systems (MCS), Reutlingen), digitized at 25,000 Hz sampling rate with 16 bit resolution using the MC_Rack software (version 4.6.2, MCS), and stored as MCD files for further offline analysis. Data from MCD files were accessed by the SPANNER software suite as well as the in-house developed in-house developed MATLAB software SSMT. For low-frequency LFP analysis, raw MEA signals sampled at 25 kHz were first downsampled to 1 kHz. For time domain analysis of local field potential oscillations, we separately determined five slowly varying time domain signal parts in the five frequency bands delta (1—4 Hz), theta (4—11 Hz), beta (11—30 Hz), gamma (30—55 Hz) and high gamma (55—100 Hz). In supplementary figure 1 A (i) to (iv), we present the FIR filter magnitude (dB) and phase (° degrees) response functions for the frequency bands delta, theta, beta and gamma. Power spectral analysis of the bandpass-filtered time domain signals is shown in supplementary figure 1 B. For frequency domain analysis of local field potential oscillations, we quantified spectral properties of the MEA signal for frequencies below 100 Hz by its power spectral density PSD (power/frequency, μV²/Hz) applying Welch’s algorithm to the 1 kHz downsampled signal. In order to simultaneously display time domain and spectral properties, we also calculated power spectrograms of the 1 kHz downsampled signal. Numerical experiments with simulated spike trains at frequencies between 1 Hz and 10 Hz superimposed on pink noise (with 1/f power spectral density) showed that these spectrogram settings permitted clear detection of onset, fundamental frequency and harmonics of the spike trains (data not shown). These spectrogram settings are thus suited to distinguish “real” electrophysiological slow LFP oscillations from artifacts created by spike trains.

## Figure legends

***Supplementary Figure 1 | Bandpass filter responses***

**(A)** Magnitude and phase responses of the used high-order linear phase finite impulse response filters for **(i)** delta, **(ii)** theta, **(iii)** beta and **(iv)** gamma frequency bands. **(B)** Exemplary Power Spectral Density graphs **(i)** for the full recorded signal and **(ii)** as overlays of the five bandpass filtered signals with colored areas for better visualization of frequency bands. The full PSD graph completely overlaps the individual bandpass filtered PSDs. Related to method section.

***Supplementary Figure 2 | Early ontogeny of synchronous bursting and oscillatory slow-wave activity in four cell lines***

**(A)** Table summarizing the used cell lines and number of experiments and replicates **(B)** Representative examples of **(i)** 2-minute raw activity traces (blue) and superimposed delta envelopes (orange) of one channel showing Type C activity in three different hiPSC lines, **(ii)** their corresponding spectrograms and **(iii)** Power Spectral Density graphs. The days after seeding on MEAs are given. Related to figure 2.

***Supplementary Figure 3 | Absence of oscillatory slow-wave activity in asynchronously active neuronal populations***

**(A, i)** Phase-contrast image, showing the morphology of a 3D-neural aggregate culture on the 9 electrodes of a 6-well micro-electrode array 6 days after seeding on the MEA plate. **(A, ii)** Corresponding spike traces (high pass filtered > 200 Hz) of the 3 × 3 electrode array, showing uncorrelated spiking activity, with **(A, iii)** zoomed-in channel 4 and spike waveform visualization. **(B, i)** Representative spectrograms of 4 individual channels and **(B, ii)** Power Spectral Density graphs showing the absence of oscillatory activity. Note that the bottom graph corresponds to the ground electrode and represents electronic 1/f noise.

***Supplementary Figure 4 | Late ontogeny of oscillatory slow-wave activity in cell line 2***

Example traces showing the oscillatory activity of hiPSC line 2, 70 days after onset of synchronous activity on MEA. **(A)** Power Spectral Density graph, where the different frequency bands are highlighted by colored boxes. **(B, i)** Representative example of 2-minute raw activity trace (blue) and superimposed delta envelope (orange) with its corresponding spectrogram. The marked rectangle is shown as zoomed-in a 6s window in **(B, ii)**. **(C)** delta, theta, beta, gamma traces and envelopes. Related to figure 4.

**Supplementary Video 1 | Spatial localization of neurons, astrocytes and oligodendroglial cells in human iPSC-3D neural aggregates.**

**Supplementary Video 2 | Oscillatory slow-wave activity depends on synaptic function and neuronal activity.**

