## Supplementary figures and images for "Ontogeny of oscillatory slow-wave and neuronal population activity in human iPSC-3D cortical circuits"

### suppl. figure 1

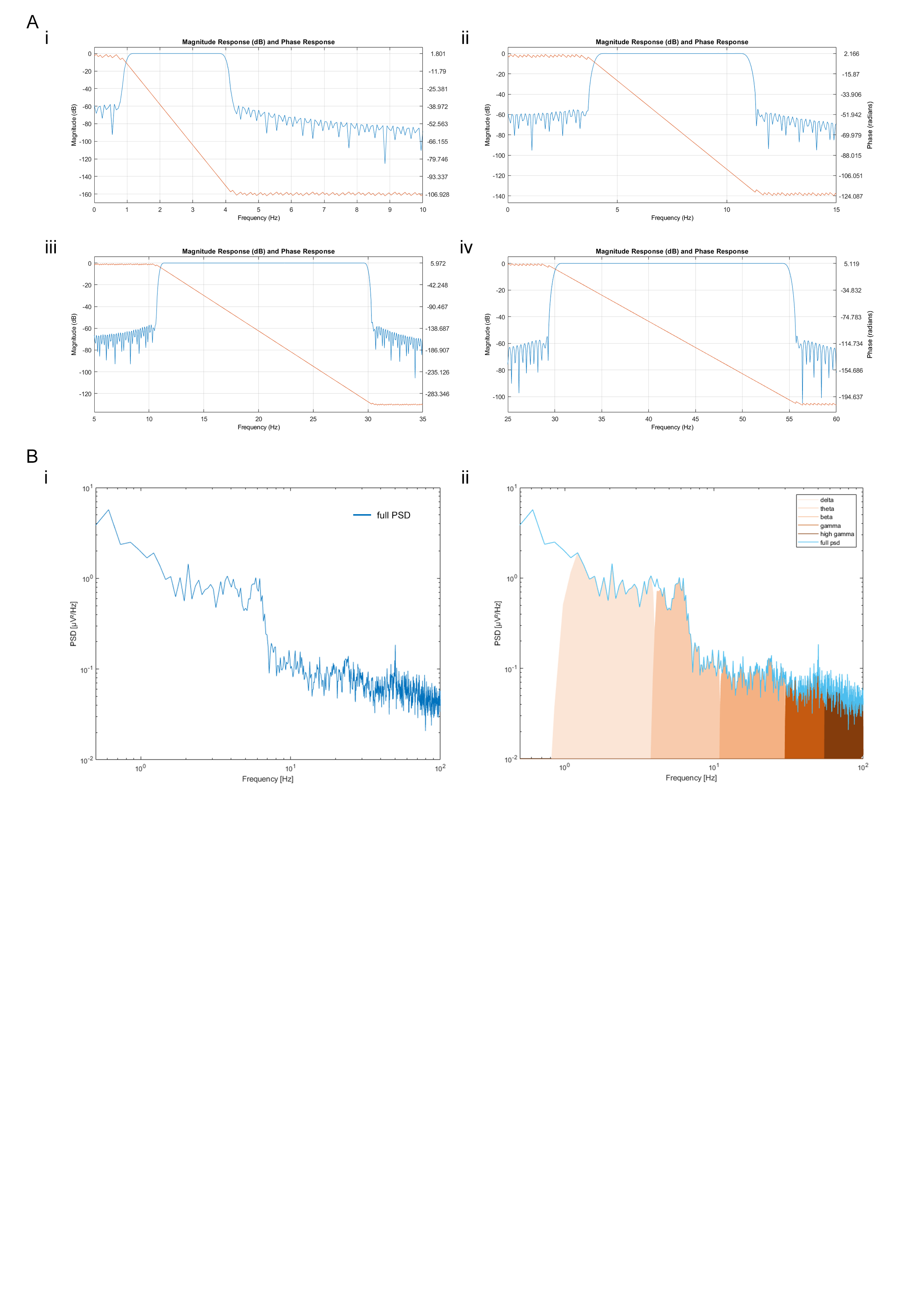

### suppl. figure 2

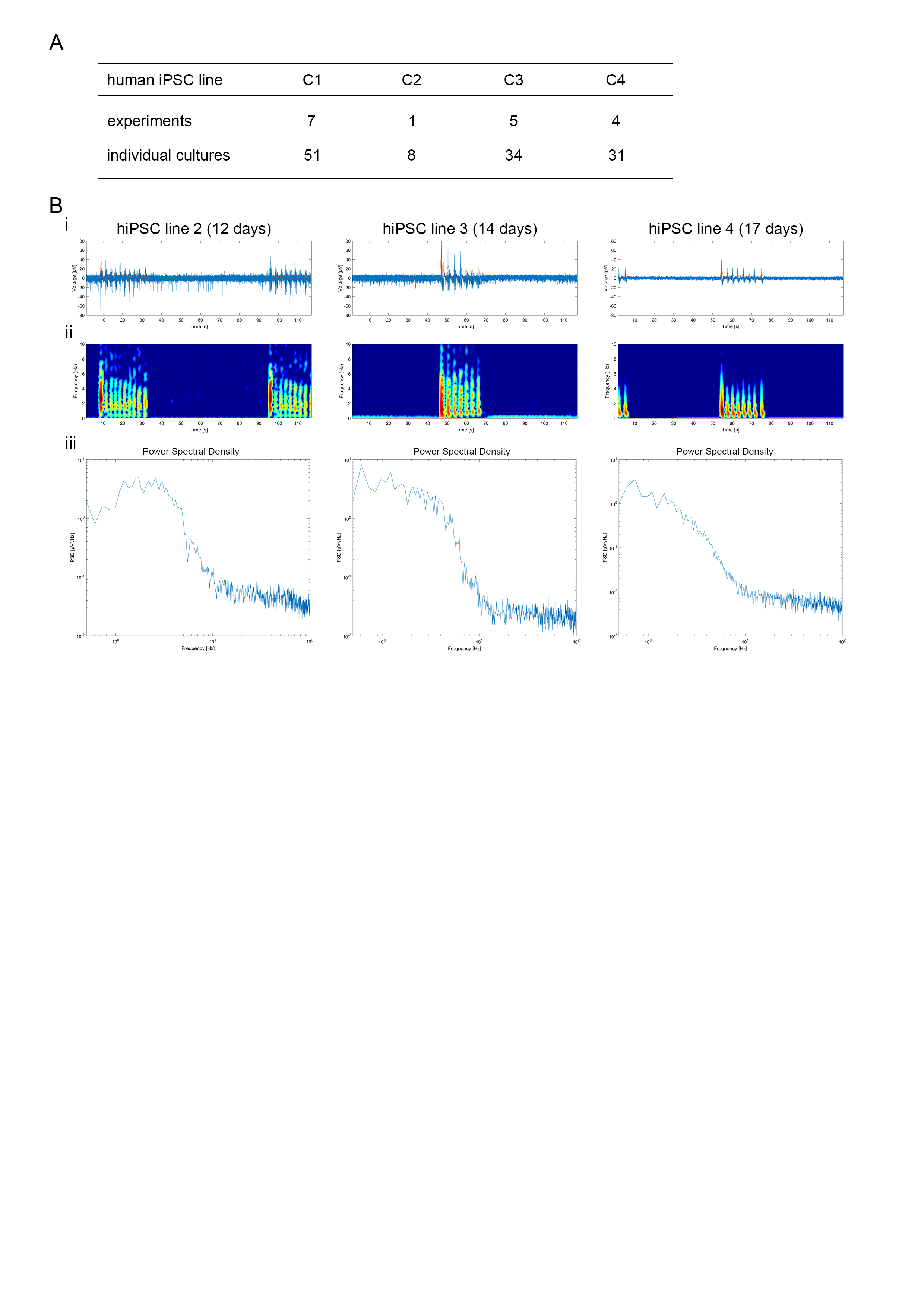

### suppl. figure 3

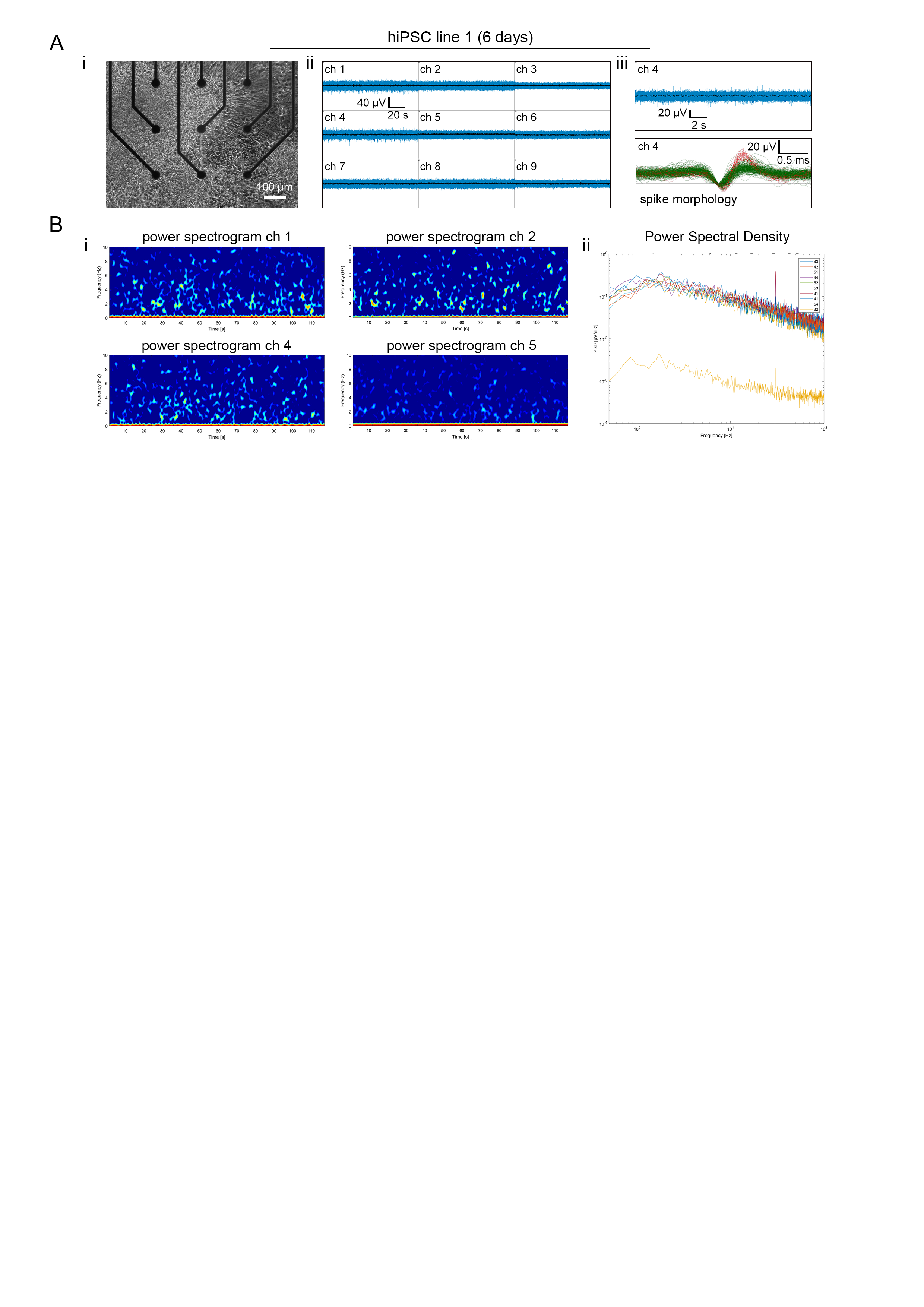

### suppl. figure 4

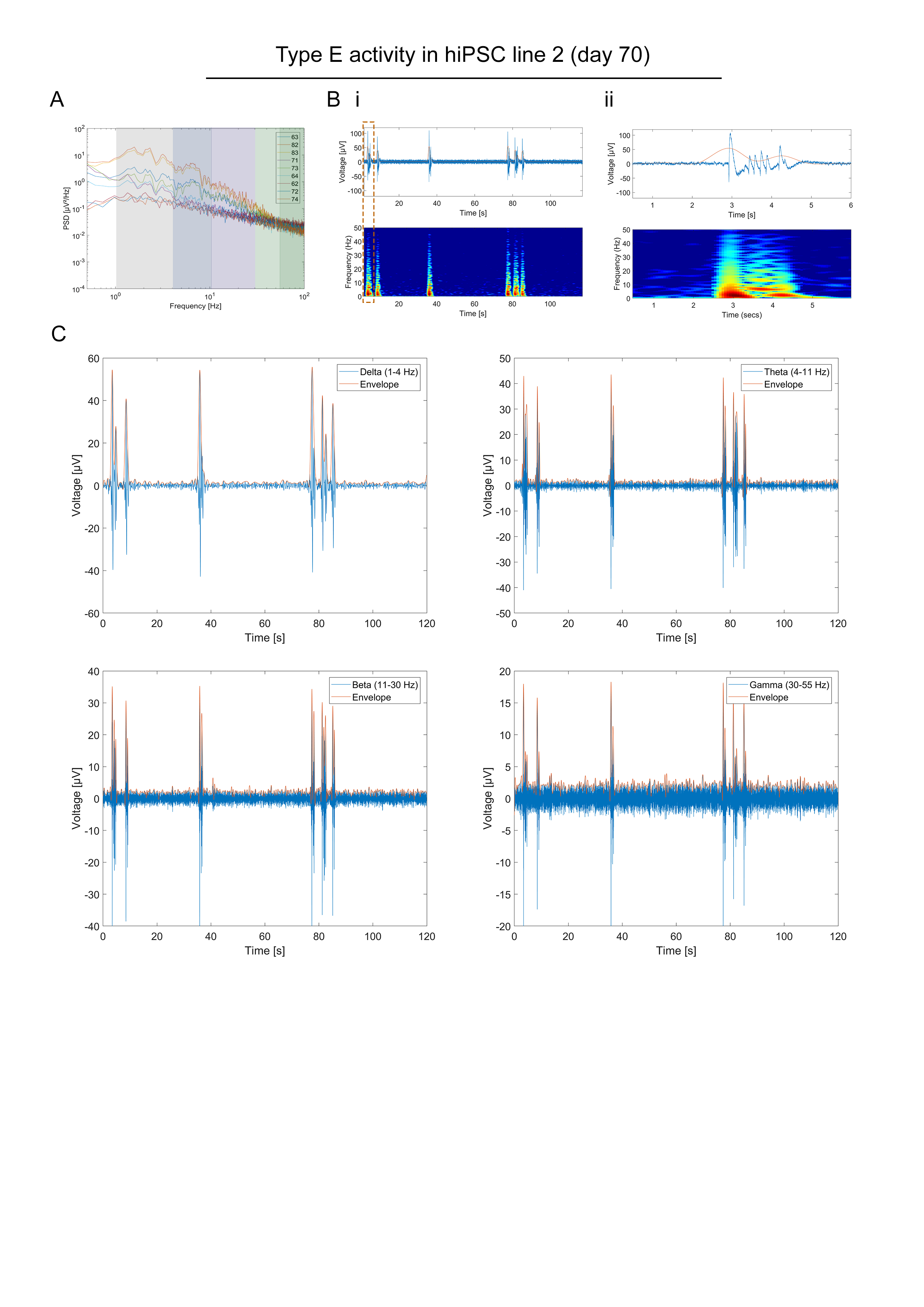
